# Leaf decomposing fungi influence *Saccharomyces paradoxus* growth across carbon environments

**DOI:** 10.1101/2023.01.06.523016

**Authors:** Samer El-Khatib, Madeleine G. Lambert, Meghan N. Reed, Melane Brito Goncalves, Primrose J. Boynton

**Affiliations:** Wheaton College, Norton, MA, USA; Georgetown University Medical Center, Washington, DC, USA; Lonza, Portsmouth, NH, USA; Rhode Island College, Providence, RI, USA

**Author notes:** Submitting and corresponding author.

## Abstract

*Saccharomyces paradoxus* is a model organism in ecology and evolution. However, its metabolism in its native habitat remains mysterious: it is frequently found growing on leaf litter, a habitat with few carbon sources that *S. paradoxus* can metabolize. We hypothesized that leaf-decomposing fungi from the same habitat break down the cellulose in leaf litter extracellularly and release glucose, supporting *S. paradoxus* growth. We found that facilitation by leaf-decomposing fungi was possible on cellulose and inhibition was common on glucose, suggesting diverse interactions between *S. paradoxus* and other fungi that have the potential to support *S. paradoxus* in nature.

## Description

Leaf litter decomposers may solve the mystery of *Saccharomyces paradoxus* metabolism in nature. *S. paradoxus* is a wild yeast and the closest described relative of *Saccharomyces cerevisiae*, the most widely researched yeast and one of the most widely researched eukaryotic model organisms (Goffeau et al. 1996; Boynton and Greig 2014). *S. paradoxus* can be found growing in leaf litter and soil close to oak trees, and less abundantly on oak tree bark (Glushakova et al. 2007; Sampaio and Gonçalves 2008; Kowallik and Greig 2016). It can grow both aerobically and anaerobically, utilizing simple sugars such as glucose as carbon sources. One common source of simple sugars in forests, honeydew (*i*.*e*., aphid excretions), has been proposed as a potential carbon source for *S. paradoxus* (Brysch-Herzberg and Seidel 2017), but surveys in New Zealand and Germany have not been able to uncover any association between *Saccharomyces* and honeydew (Serjeant et al. 2008; Kowallik 2015). *S. paradoxus* also cannot metabolize cellulose, one of the most common polymers in leaf litter (Van Rensburg et al. 1998; Dashko et al. 2014). The primary component of the plant leaf cell walls in leaf litter is cellulose, and glucose and other simple monosaccharides are generally scarce on leaf litter (Sampaio and Gonçalves 2008; Keegstra 2010). It is unclear how *S. paradoxus* survives and reproduces in such cellulose-rich and glucose-poor environments: here, we investigate how interactions with nearby fungi might facilitate *S. paradoxus* metabolism.

Diverse leaf fungal communities are the main decomposers of leaf litter, playing an essential role globally in plant polymer decomposition and the carbon cycle (Kjøller and Struwe 1982; Benocci et al. 2017). Fungi that occupy the same habitat as *S. paradoxus* decompose leaf litter through extracellular cellulolytic enzymes, such as cellulases and glucanases (Krishna and Mohan 2017). Previous research has shown that nearby microorganisms can facilitate or inhibit *S. paradoxus* growth on oak bark (Kowallik et al. 2015). However, the mechanisms behind these interactions remain unknown. We hypothesize that cellulose-degrading fungi facilitate the survival of *S. paradoxus* in leaf litter through extracellular breakdown of leaf cellulose and the consequent release of more accessible sugars such as glucose. Conversely, we hypothesize that cellulose-degrading fungi might consume free glucose, a carbon source otherwise conducive to *S. paradoxus* growth, and inhibit *S. paradoxus* when cellulose is not available.

To test our hypotheses, we first isolated five *S. paradoxus* strains and five leaf-degrading fungal isolates with diverse morphologies (**Figure 1a-b**) from partially decomposed oak leaves in a New England forest. The leaf-degrading fungi were identified using ribosomal DNA sequences as members of the genera *Trichoderma, Biscogniauxia, Coniochaeta, Chaetomium*, and *Nemania*. These genera include characterized leaf endophytes, leaf litter decomposers, and wood decomposing fungi (Nugent et al. 2005; Fukasawa et al. 2011; Osono et al. 2011; Helaly et al. 2018; Mäkelä et al. 2021). We confirmed that all five litter fungi, but not *S. paradoxus*, can grow on a growth medium containing cellulose as the sole carbon source, and then further explored the interactions between *S. paradoxus* and the other fungi in different carbon environments.

**Figure 1:**
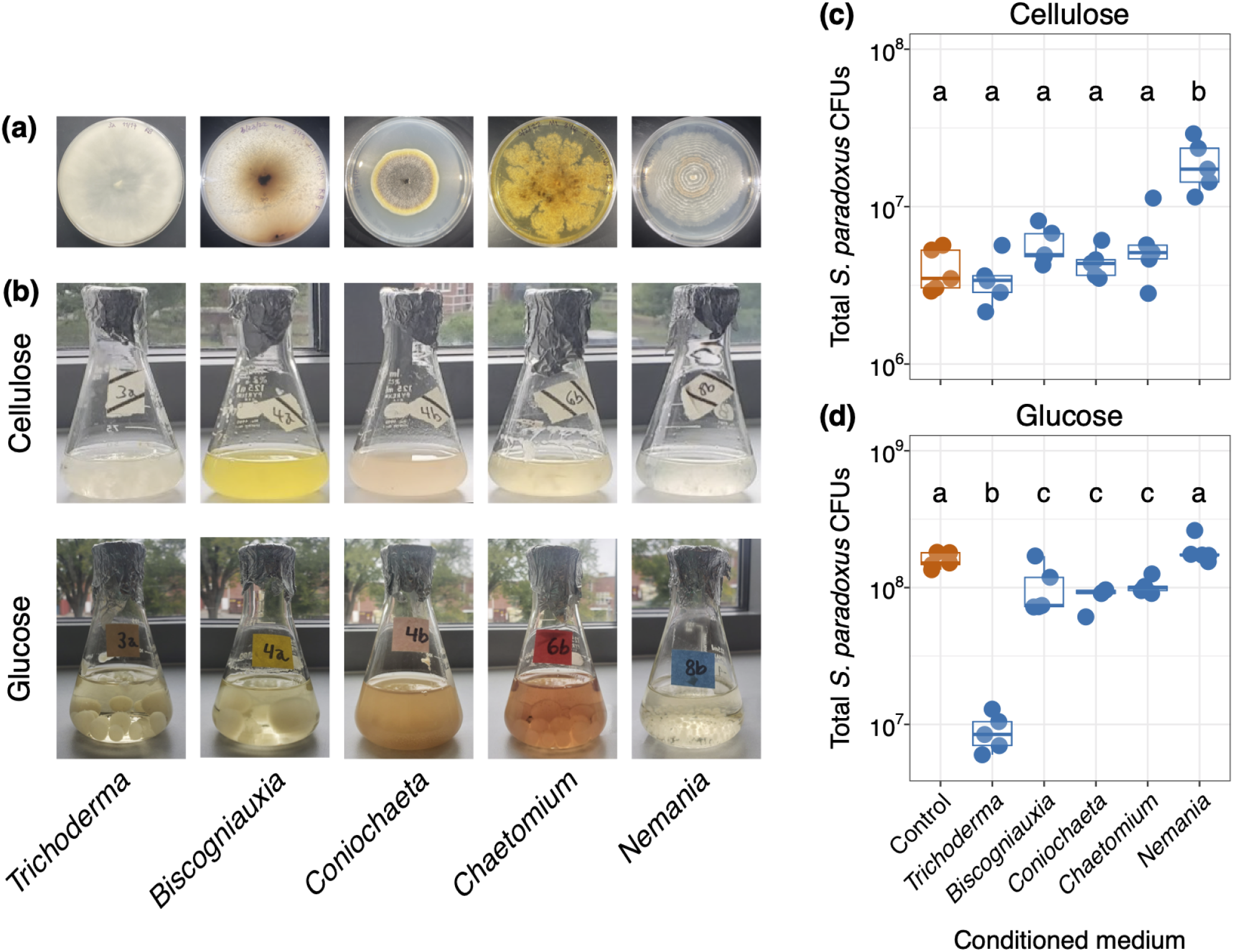
Leaf litter decomposing fungi and their interactions with *S. paradoxus*. **(a)** Five filamentous fungi isolated from forest oak leaf litter growing on solid glucose medium. **(b)** The five fungi in flasks of liquid cellulose (top) and liquid glucose (bottom) media. **(c)** The total number of *S. paradoxus* colony forming units (CFUs) grown after 26 hours in 5 ml cellulose medium conditioned by each of five litter-decomposing fungi, or in control unaltered media (five replicate *S. paradoxus* strains per treatment; each *S. paradoxus* strain is represented by a single dot per treatment). Conditioned media were prepared after 96 hours of fungal growth in liquid media, followed by filter-sterilization of used media. *Nemania* increased the number of *S. paradoxus* CFUs to 4.7 times the control (one-way ANOVA adjusted R^2^ = 0.72, F = 15.6, degrees of freedom = 5,24, p < 10^−6^). In contrast, *S. paradoxus* produced equivalent numbers of CFUs in the unaltered control medium as in media conditioned by *Trichoderma, Biscogniauxia, Coniochaeta*, and *Chaetomium*. **(d)** The total number of *S. paradoxus* CFUs grown in glucose medium conditioned by each of the five litter-decomposing fungi, plus an untreated control. *S. paradoxus* produced considerably fewer CFUs after conditioning by *Trichoderma* (0.06 times the number of CFUs produced in untreated glucose medium; one-way ANOVA adjusted R^2^ = 0.81, F = 25.69, degrees of freedom = 5,24, p < 10^−8^). *S. paradoxus* also produced fewer CFUs after conditioning by *Biscogniauxia, Coniochaeta, and Chaetomium* than in unaltered glucose medium (0.61 times the control number of CFUs). *Nemania* did not significantly change the mean number of *S. paradoxus* CFUs compared to the control. The y-axes are presented on a log_10_ scale in figures 1c and 1d, and different letters in each of figures 1c and 1d indicate significantly different means (Tukey post-hoc HSD p < 0.05).

We carried out spent-media experiments to test our hypothesis that the leaf litter decomposing fungi would facilitate *S. paradoxus* growth on cellulose by breaking down the polymer into simpler sugars. We first allowed each fungus to grow on cellulose medium and to change the environment, including presumably through the release of cellulolytic enzymes or metabolites into the medium. Cellulose medium, used for spent-media experiments, differs from cellulose screening medium, used to confirm growth with cellulose as a sole carbon source, in that it contains peptone and can support some *S. paradoxus* growth. We then filtered out each fungus and grew *S. paradoxus* on the used medium, quantifying *S. paradoxus* growth using colony-forming unit (CFU) counts for each fungal treatment after 26 hours. One of the five fungi, *Nemania*, increased the mean number of *S. paradoxus* CFUs produced to 4.7 times the control untreated medium, but the other four fungi had no significant effect on growth (**Figure 1c**, ANOVA adjusted R^2^ = 0.72, F = 15.6, degrees of freedom = 24,5, p < 10^−6^; Tukey post-hoc comparison between the control and *Nemania* treatments p < 10^−5^).

To further investigate the effects that leaf litter decomposing fungi have on *S. paradoxus* growth, we repeated the experiment with glucose medium, an environment permissive to *S. paradoxus* growth. We hypothesized that the litter decomposing fungi would use up free glucose, inhibiting *S. paradoxus* growth. In glucose medium, the previously identified facilitator, *Nemania*, had no significant effect on *S. paradoxus* growth, while *Trichoderma* had an extremely inhibitory effect, decreasing the mean number of *S. paradoxus* CFUs to 0.06 times the control. *Biscogniauxia, Coniochaeta*, and *Chaetomium* all also inhibited growth compared to the control but to a lesser extent, decreasing the mean number of CFUs to 0.61 times the control (**Figure 1d**, ANOVA adjusted R^2^ = 0.8098, F = 25.69, degrees of freedom = 24,5, p < 10^−8^; Tukey post-hoc p = 10^−7^ for the comparison between *the Trichoderma* and the control, and p ≤ 0.036 for comparisons between each of *Biscogniauxia, Coniochaeta*, and *Chaetomium* and control). The varying degrees of inhibition by different fungi on glucose indicates that certain leaf litter species are more efficiently antagonistic to *S. paradoxus* than others.

Our results partially support our hypotheses and highlight the diversity of possible interactions between *S. paradoxus* and other litter decomposing fungi. *S. paradoxus* growth on cellulose was facilitated by at least one litter decomposing fungus, while growth on glucose was inhibited to varying degrees by four of the fungi. *Nemania* likely facilitates *S. paradoxus* on cellulose by releasing cellulolytic enzymes and enabling opportunistic *S. paradoxus* glucose consumption after the cellulose is broken down. *Nemania*’s lack of inhibition in glucose could also indicate that *Nemania* doesn’t consume glucose quickly enough to prevent *S. paradoxus* from accessing it, lending a rare but well-timed opportunity for *S. paradoxus* to grow in unfavorable carbon environments such as leaf litter.

Other interactions between *S. paradoxus* and leaf decomposing fungi (**Figure 1c-d**) may be due to more complex interactions, possibly involving primary metabolism or secreted antagonistic molecules. We speculate that cellulose-degrading fungi might be especially effective at removing glucose from the environment when cellulolytic enzymes are not needed. High concentrations of glucose repress cellulolytic enzyme production (Strauss et al. 1995; Suto and Tomita, 2001), allowing more resources to be devoted to other cell functions, perhaps including glucose uptake. This increased glucose uptake and removal from the environment may allow litter decomposing fungi to more effectively inhibit *S. paradoxus* in glucose than cellulose (**Figures 1c-d**). Litter decomposing fungi might also release metabolites that could directly inhibit *S. paradoxus*. Qualitative observations of differences in pigmentation among the fungi in cellulose and glucose growth media (**Figure 1b**) support this mechanism. Additionally, fungi in the *Sordariomycetes* class, to which *Chaetomium, Coniochaeta*, and *Trichoderma* belong, and others in the *Xylariales* order, to which *Biscogniauxia* and *Nemania* belong, release active secondary metabolites, including pigments (Stamps et al. 2015; Bills and Gloer, 2016; Helaly et al. 2018; Becker and Stadler, 2021), with some fungi doing so when associated with other organisms (Lamacchia et al. 2016). In this case, fungi growing in glucose might be able to expend more energy towards the production of active secondary antagonistic metabolites because cellulolytic enzyme production is not necessary.

*S. paradoxus* growth on different carbon sources is contingent on interactions among *S. paradoxus*, leaf litter fungi, and the environment. Litter decomposers can create conditions conducive to *S. paradoxus*, or they can inhibit *S. paradoxus*, depending on the nutrient environment. *S. paradoxus* growth in its natural cellulose environment most likely depends on the presence of facilitator fungi such as *Nemania*, the lack of antagonistic species such as *Trichoderma*, and well-timed opportunistic glucose consumption. Future research will explore the roles of cellulases and secondary metabolites in interactions between litter decomposing fungi and *S. paradoxus* in nature.

## Methods

### Isolation and identification of *S. paradoxus* and leaf litter decomposing fungi

All fungi were isolated from oak leaf litter in a mixed temperate forest in Norton, MA, USA (coordinates 41.96, -71.18). Five *S. paradoxus* strains were isolated from leaf litter in the summer of 2021 as in Boynton et al. (2019). First, 134 yeast colonies were isolated by directly plating soil directly underneath leaf litter onto the isolation medium PIM1 (Sniegowski et al. 2002; Boynton et al., 2019). All media recipes are provided in the reagents section below. Colonies were then screened for *Saccharomyces*-like spore morphology, and *S. paradoxus* colonies were identified using *S. paradoxus* specific PCR primers, as in Robinson et al. (2016), or by sequencing the ribosomal internal transcribed spacer region using primers ITS1 and ITS4 (White et al. 1990).

Filamentous fungi were isolated from leaf litter next to three oak trees; partially degraded fallen leaves (edges were degraded but not the center) were collected in the fall of 2021, aseptically contained in sterile plastic bags, and stored at approximately 4°C for up to 24 hours before fungal isolation. Seven-millimeter diameter leaf discs were punched out from each leaf three times with a sterile hole punch. Discs were surface sterilized as in Osono & Takeda (2001) and were arranged on filamentous fungi isolation medium and incubated at 28°C until fungal mycelium could be seen growing from leaf cut edges. Five fungi were subcultured onto glucose medium and pure cultures were maintained on glucose medium for the duration of the study. Growth on cellulose as a sole carbon source was confirmed by observing growth on cellulose screening medium. The fungi were identified by sequencing portions of large ribosomal subunit and ribosomal internal transcribed spacer DNA using primers detailed in the reagents section below. Fungi were identified to genera using NCBI BLAST (Altschul et al. 1990; Sayers et al. 2022) and querying databases associated with 26S RNA and internal transcribed spacer regions from Fungi type and reference material. Internal transcribed spacer and partial 26S RNA sequences were deposited in GenBank (Accession numbers OQ145425-OQ145429, OQ145437-145441).

### Media conditioning experiments

To understand the impacts of each leaf litter fungus on *S. paradoxus*, we grew *S. paradoxus* on cellulose and glucose media that had first been conditioned by each leaf litter fungus. Each of the leaf litter fungi was inoculated into a flask containing 75 mL of liquid medium. Flasks were incubated at 28°C for 96 hours with 140 rpm shaking. After growth, the conditioned medium from each flask was filtered through a 0.2 µm filter, dispensed into 5 tubes (5 ml conditioned medium per tube), and each tube was inoculated with one of five *S. paradoxus* strains by transferring colonies into each tube from a petri dish using a sterile wooden stick. Tubes were incubated at 28°C with 140 rpm shaking for 26 hours, after which *S. paradoxus* colony forming units (CFUs) were counted on YEPD solid medium. In untreated cellulose medium, *S. paradoxus* could grow a little bit, most likely by using peptone in the medium as a carbon source. One-way ANOVAs with Tukey’s HSD post-hoc tests were performed for each medium with the conditioning fungus as a predictor and number of *S. paradoxus* cells produced as a response variable. Statistical tests were performed using R version 4.1.2, with plots produced using the ggplot2 package (Wickham 2016; R Core Team, 2021).

## Reagents

**Fungal strains**

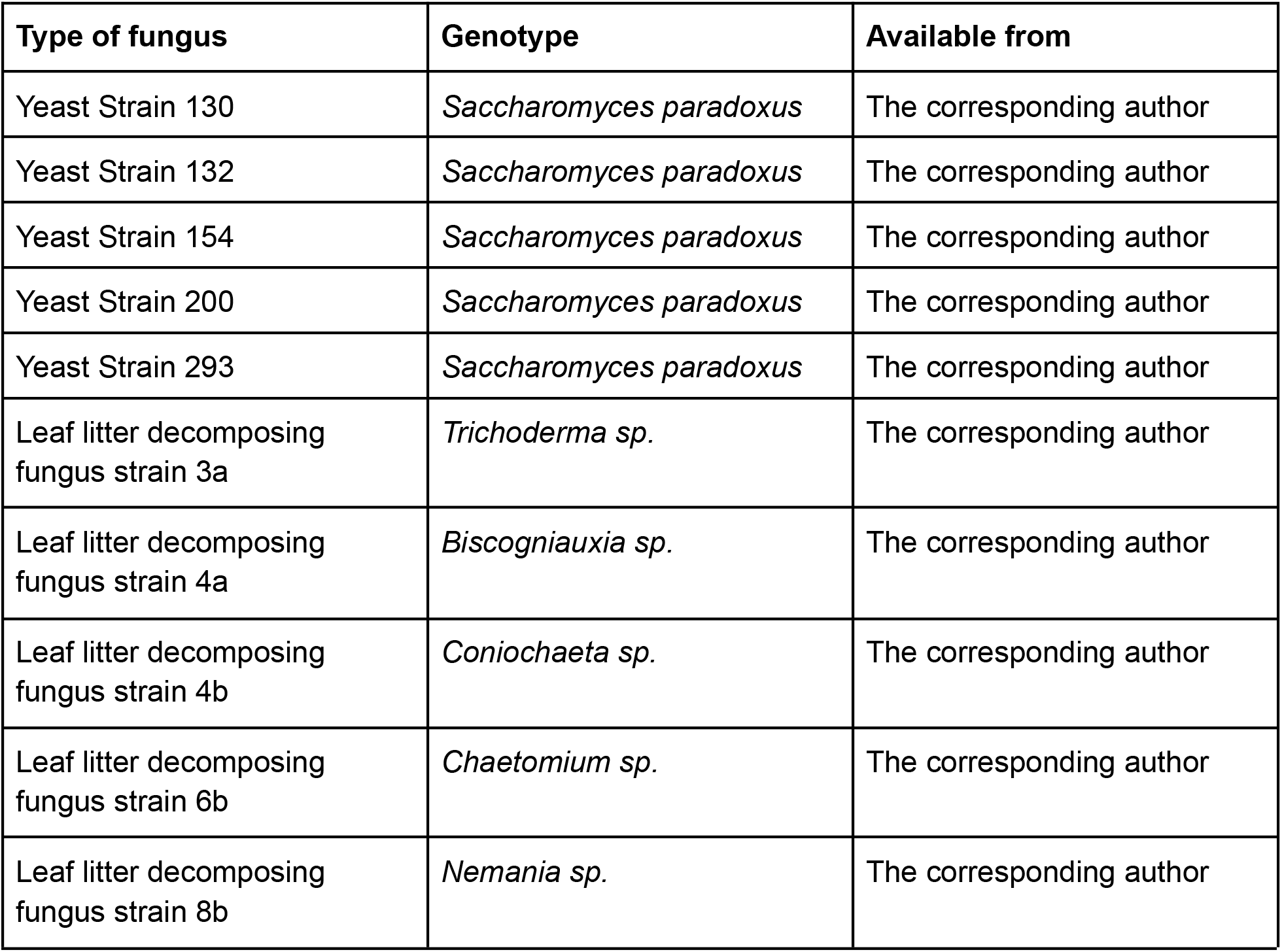

**Primers**

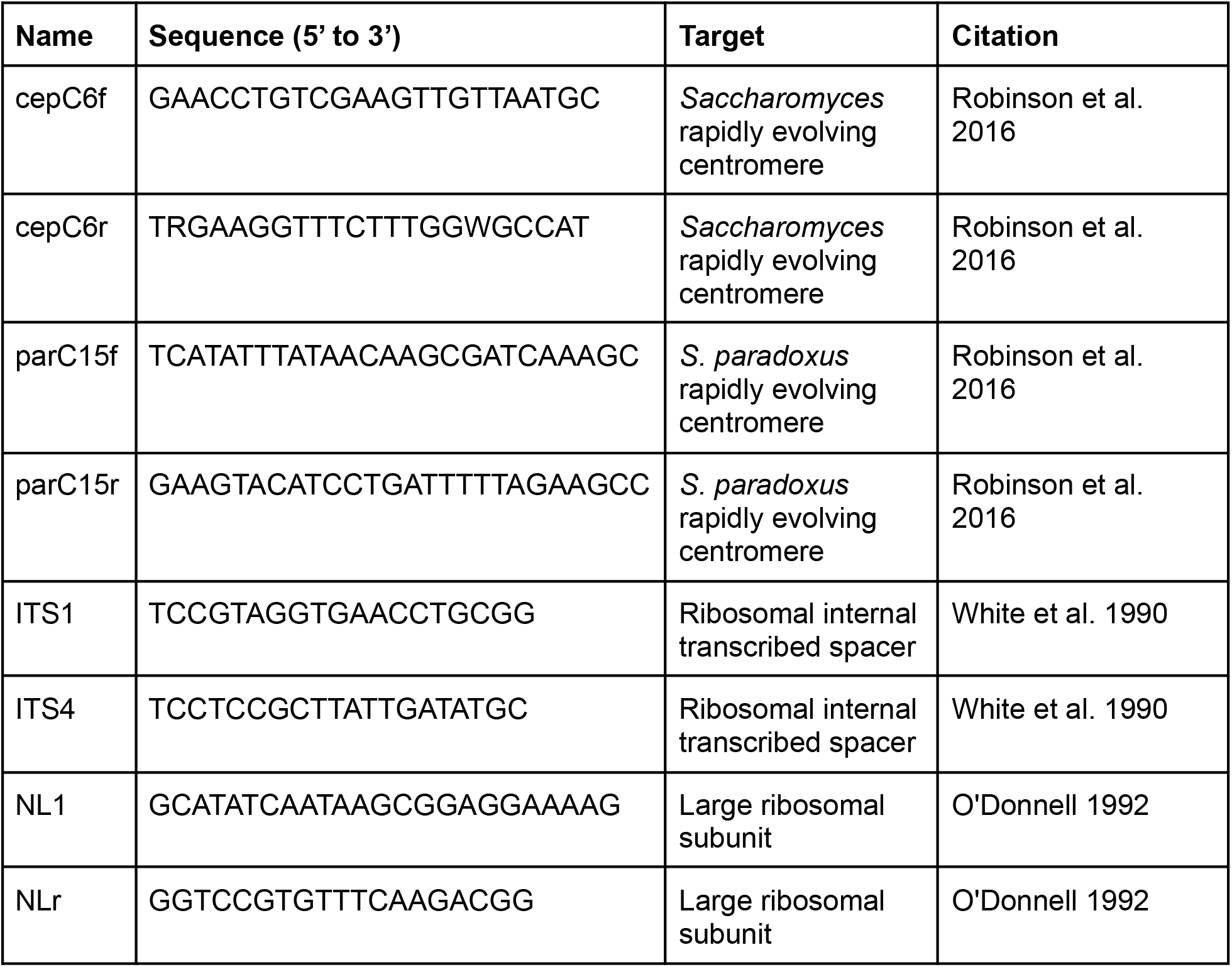

**Media**

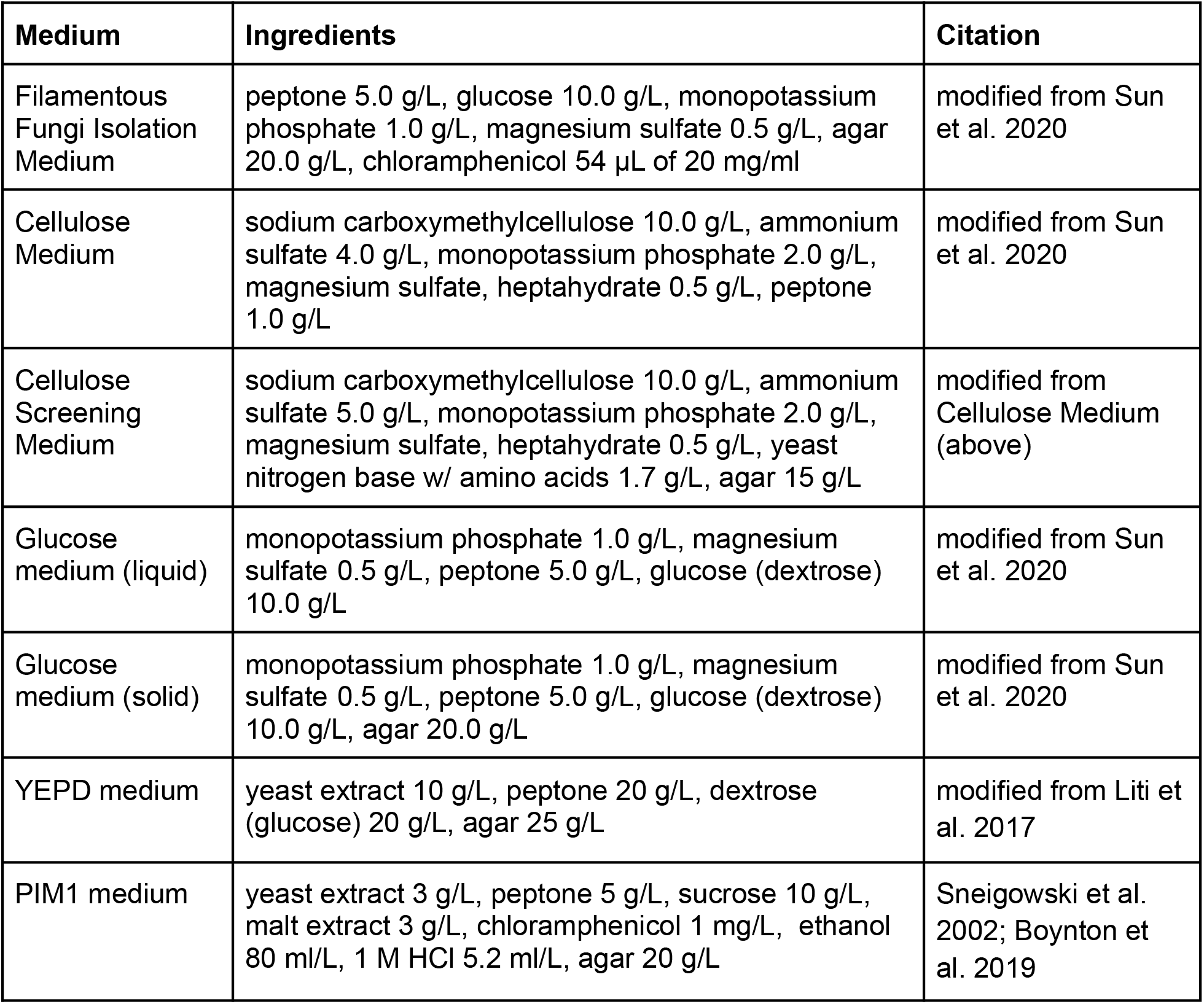

## Extended Data

File name: Saccharomyces paradoxus conditioning media experiment.csv

Description: Data table (.csv format) of treatments and CFUs for Figure 1c-d.

## Author contributions

Samer El-Khatib: Data curation, formal analysis, funding acquisition, investigation, methodology, visualization, writing–original draft, writing–review & editing Madeleine G. Lambert: Conceptualization, data curation, formal analysis, funding acquisition, investigation, methodology, visualization, writing–review & editing Meghan N. Reed: Funding acquisition, investigation, methodology, writing–review & editing Melane Bitro Goncalves: Funding acquisition, investigation, writing–review & editing Primrose J. Boynton: Conceptualization, data curation, formal analysis, funding acquisition, methodology, project administration, supervision, validation, visualization, writing–original draft, writing–review & editing

